# Cryo-EM reconstruction of AlfA from *Bacillus subtilis* reveals the structure of a simplified actin-like filament at 3.4 Å resolution

**DOI:** 10.1101/190744

**Authors:** Andrzej Szewczak-Harris, Jan Löwe

## Abstract

Low copy-number plasmid pLS32 of *Bacillus subtilis* subsp. *natto* contains a partitioning system that ensures segregation of plasmid copies during cell division. The partitioning locus comprises actin-like protein AlfA, adaptor protein AlfB and the centromeric sequence *parN*. Similar to the ParMRC partitioning system from *E. coli* plasmid R1, AlfA filaments form actin-like double helical filaments that arrange into an antiparallel bipolar spindle, which attaches its growing ends to sister plasmids, through interactions with AlfB and *parN*. Since, compared with ParM and other actin-like proteins, AlfA is highly diverged in sequence, we determined the atomic structure of non-bundling AlfA filaments to 3.4 Å resolution by cryo-EM. The structure reveals how the deletion of subdomain IIB of the canonical actin-fold has been accommodated by unique longitudinal and lateral contacts, whilst still enabling formation of left-handed, double helical, polar and staggered filaments that are architecturally similar to ParM. Through cryo-EM reconstruction of bundling AlfA filaments we obtained a pseudo-atomic model of AlfA doublets: the assembly of two filaments. The filaments are antiparallel, as required by the segregation mechanism, and exactly anti-phasic with 8-fold integer helical symmetry, to enable efficient doublet formation. The structure of AlfA filaments and doublets shows, in atomic detail, signs of the strong evolutionary pressure for simplicity, placed on plasmids: deletion of an entire domain of the actin fold is compensated by changes to all interfaces so that the required properties of polymerisation, nucleotide hydrolysis and antiparallel doublet formation are retained to fulfil the system's biological raison d'être.

**Significance Statement:** Protein filaments perform a vast array of functions inside almost all living cells. Actin-like proteins in archaea and bacteria have previously been found to form a surprising diversity of filament architectures, reflecting their divergent cellular roles. Actin-like AlfA is unique in that it is much smaller than all other filament forming actin-like proteins. With an atomic structure of the AlfA filament, obtained by high-resolution electron cryo-microscopy, we have revealed—at atomic level of detail—how AlfA filaments form dynamic filaments capable of transporting plasmid DNA in cells and how these filaments arrange into antiparallel bundles required for the segregation mechanism.

## Introduction

Actin-like proteins are defined by structural and functional homology to actin, the first identified member of the family and the fundamental building block of the eukaryotic microfilament cytoskeleton. Actin-like proteins make up a functionally diverse and evolutionarily near-ubiquitous family of proteins. A unifying feature of actin-like proteins is the ability to form filaments *in vitro* and *in vivo*. All studied actin-like proteins bind and catalyse the hydrolysis of nucleotide triphosphates, especially ATP, processes which respectively promote filament growth and subsequent disassembly (1). Sequence database searches and structural studies revealed the presence of numerous actin homologues in bacteria and archaea, helping to establish the notion of the prokaryotic origin of actin-like cytoskeletal proteins (2). The roles that actins fulfil in cells reflect their ability to polymerise and are especially diverse in prokaryotes, where they function in cytokinesis, morphogenesis and DNA replication. Amongst the chromosomally encoded bacterial and archaeal actin-like proteins, we find: FtsA, an element of the cytokinetic (Z-) ring (3), MreB, a membrane-binding protein involved in the organisation of the cell wall synthesis and thus cell shape maintenance (2), as well as MamK, which organises the distribution of magnetosomes, iron-containing organelles found in magnetotactic bacteria (4). Within the archaeal phylum Crenarcheota, actin-like protein crenactin has been identified, and has been shown to be the closest so far in filament architecture and monomer fold to eukaryotic actin. The function of crenactin remains to be elucidated (5). In addition to the housekeeping roles described above, a separate functional group evolved in the actin-like family to aid the replication of certain low copy-number plasmids, where actin-like filaments enable efficient segregation of plasmid DNA.

As plasmid DNA is replicated separately from chromosomal DNA, mechanisms that ensure the carry-over of genomic material to the progeny cells do not apply to plasmids. This issue has more serious consequences for low copy-number plasmids, and in order to address it some of them evolved mechanisms that employ filamentous proteins to physically separate two plasmid copies and move them across cells (6). The best studied of these cases involves ParM, an actin-like protein, encoded by the *parMRC* locus of the R1 plasmid in Gram-negative *Escherichia coli*. The locus encodes for the tripartite ParMRC system that, besides ParM, contains the DNA-binding protein ParR and a centromeric sequence *parC* (7). Principles governing plasmid segregation by the ParMRC system are well understood mechanistically. Atomic structures of the ParM filament and monomer in various nucleotide states (8-10), as well as with ParR peptide (9), and ParR bound to *parC* (11), have been solved with X-ray crystallography and electron cryo-microscopy (cryo-EM). Biochemical reconstitution assays and total internal reflection (TIRF) microscopy have complemented these studies to reveal the dynamics of the filament assembly and DNA segregation (9). According to the current model, ParR binds to the *parC* sequence and acts as a seed for polymerisation of ParM filaments after replication of the *parC* centromere-like regions. Two filaments, each attached to a sister copy of the plasmid DNA, bundle in antiparallel pairs and grow through addition of subunits at the tips, pushing the plasmids apart (10). Conceptually, the ParMRC system can be regarded as a truly minimalistic mitotic spindle apparatus.

A similar system was identified in a strain of Gram-positive bacterium *Bacillus subtilis*, subsp. *natto*, used in the fermentation of traditional Japanese food *nattō* from soya bean (*Glycine max*). There, a cryptic, low copy-number plasmid pLS32 was found to contain a region responsible for conferring plasmid stability during sporulation and cell replication (12-14). Analogous to ParMRC, the plasmid-maintenance system of pLS32, contains an actin-like protein AlfA (15), a DNA-binding adaptor AlfB of unknown three-dimensional structure, and a centromeric sequence *parN* (16). Although the dynamics of the filament assembly and disassembly differ between AlfA and ParM, the basic principles of plasmid segregation are preserved: AlfA filaments, bound to the plasmid’s *parN* sequence via the adaptor AlfB, bundle together to form antiparallel and bipolar spindles that move DNA apart (17, 18). From the structural point of view, the system remains less well characterised than ParMRC, as the atomic details of the AlfA and AlfB proteins have not yet been elucidated. As indicated by sequence alignments, the 31-kDa AlfA (275 amino acid long), is so far the smallest of the identified actin-like proteins. Its reduced size is a consequence of the deletion of one (IIB) of the four subdomains characteristic of all other members of the family. If confirmed by an atomic structural model, the deletion poses interesting questions regarding the maintenance of filament functionality in light of the known structures of other actin-like filaments, including actin itself and ParM. In order to study how the effects of the deletion are mitigated by structural adaptations in other parts of the protein, we decided to reconstruct the atomic structure of AlfA filament using electron cryomicroscopy (cryo-EM).

## Results and Discussion

### Cryo-EM reconstruction of AlfA shows double-helical, parallel, left-handed filaments at near-atomic resolution

A previous study established that AlfA polymerised in the presence of ATP to form doublets of protofilaments (strands) that wrap around each other in a left-handed helix (18). The analysis was carried out with conventional, room temperature EM using negatively stained samples, which limited the resolution to around 15 Å. To obtain a detailed, near-atomic model of the filament structure, we turned to cryo-EM, which has recently shown much success in providing such information for actin, ParM, MamK and crenactin filaments (4, 5, 10, 19).

We began our study of the AlfA filament structure with the wild-type protein sequence derived from the pLS32 plasmid, which was codon-optimised, synthesised and cloned into a vector for expression in *E. coli*. Purified protein was polymerised via the addition of ATP, deposited on a cryo-EM sample grid, vitrified and imaged in an electron microscope (Fig. S1A). Similar to previously published negative-stain images of native AlfA (18), we observed heavy bundling of the filaments, which was alleviated only by raising salt (KCl) concentrations to 1000 mM (Fig. S1B-D). Disruption of bundles and obtaining single AlfA filaments is necessary for the purpose of filament reconstruction; however, increasing salt concentration significantly decreases contrast of the cryo-EM images (Fig. S1B-D). This, in effect, leads to a decrease in signal-to-noise ratio, preventing high-resolution structure determination. In order to obtain single filaments of AlfA, we turned to a non-bundling protein mutant, described previously and used for dynamic studies of the plasmid’s segregation system (18). The non-bundling AlfA is a quadruple mutant (K21A, K22A, K101A, K102A), designed to abolish the electrostatic interactions between lysine and glutamate residues that have been implicated, through structural modelling, in AlfA bundle formation. Cryo-EM imaging of polymerised non-bundling AlfA mutant confirmed the presence of single filaments in a low-salt background suitable for further image processing and structure solution (Fig. S1E).

The procedure of structure solution was carried out exploiting the helical reconstruction capabilities implemented in RELION (20). Briefly, the method involves single particle-like processing of helical assemblies in a Bayesian, empirical framework, which uses a marginalised likelihood function and a prior on the reconstruction to dampen high spatial-frequency information, where there is an absence of relevant empirical data (21). Initial, reference-free 2D classification of the AlfA sample showed that individual filament segments could be effectively averaged to reveal the double-helical nature of the filament with staggered subunits as well as secondary structure features (Fig. S1F). As 3D reference for the 3D refinement we used a ParM crystal structure (PDB: 1MWM), which was extended *in silico* into a helix using parameters deduced from the 2D classes and previously published data (18). The model was low-pass filtered to 30 Å to prevent emergence of higher-resolution structural features not present in the experimental data. The procedure involved real-space optimisation of helical symmetry parameters at each of the cycles of the refinement and yielded final values of 156.5° (twist) and 24.9 Å (rise). In order to prevent overfitting, overlapping segments from each filament were kept in the same half-set that was used to calculate Fourier shell correlation (FSC).

The cryo-EM map obtained in the final step of the 3D refinement has an overall resolution of 3.4 Å (Fig. 1B-C), calculated using the FSC0.143 gold-standard criterion (22). As expected at this level of detail, the map clearly resolves secondary structure features as well as side chain densities for the bulkier residues (Fig. 1E-G). The density of the nucleotide bound in the catalytic pocket is clear and allowed the fitting of the hydrolysed nucleotide ADP, as well as the co-ordinating magnesium ion—which is also well resolved (Fig. 1H). Based on the density map, a complete atomic model of AlfA monomer (Fig. 1D, see next section) could be built and refined. Expanding the model with the helical symmetry parameters determined during the 3D refinement allowed us to recreate the whole filament's atomic structure (Fig. 1A). The model shows a left-handed, double-helical, parallel (polar) filament with staggered subunits and each subunit bound to ADP, as expressed by the formula 2p(AlfA_AXP_)^N^ (23). The AlfA filament is similar to the ParM filament in terms of its order, polarity, and handedness, but forms tighter coils, as reflected in the difference of the helical twist: 156.5° for AlfA (left turn of 47° between subunits along one AlfA protofilament) and 165.1° for ParM (left turn of 29.8° between subunits) (Fig. S2).

**Fig. 1.**
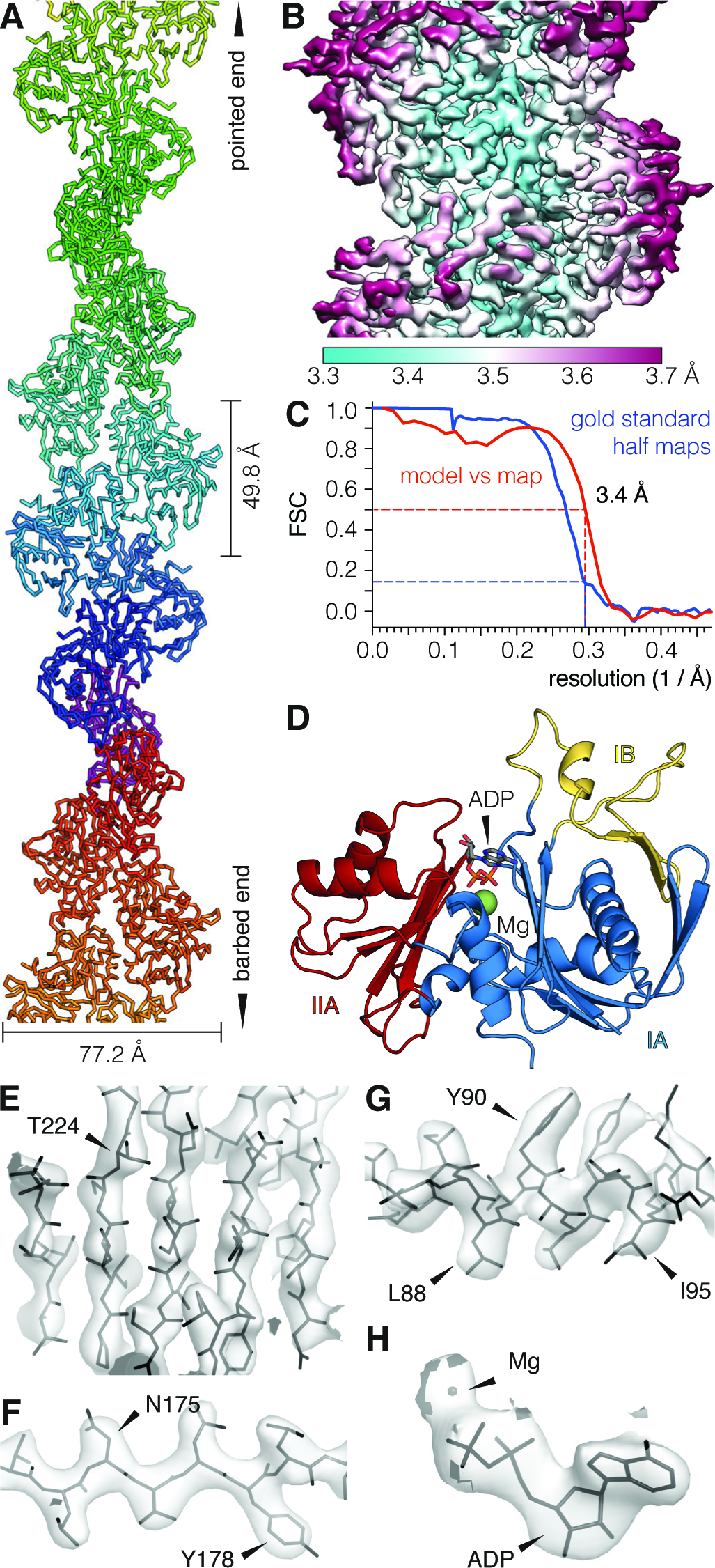
Cryo-EM reconstruction of AlfA filament. (*A*) Refined atomic model of AlfA filament in ribbon representation. The subunits are staggered in a left-handed, double-helical, parallel (polar) filament, with a rise of 24.9 Å and a 156.5° twist. (*B*) Central portion of the cryo-EM density used to build the atomic model of AlfA, coloured according to local resolution of the map. (*C*) Gold-standard FSC curves as calculated in RELION (20), showing the correlation between the half-datasets (blue line) as well as between the map and the model (red line). The overall resolution is 3.4 Å, as judged by the FSC0.143 and FSC0.5 criteria for the two correlations (blue and red dashed lines). (*D*) Atomic model of AlfA monomer, in cartoon representation, built from and refined against the cryo-EM density. Highlighted, are subdomains IB, IA and IIA of the actin fold (yellow, blue and red, respectively), as well as ADP nucleotide. (*E-H*) Selected portions of the cryo-EM density, showing the reconstructed map around representative secondary structural features: β-sheet (*E*), strand (*F*), and α-helix (*G*), as well as nucleotide density with co-ordinated magnesium ion (*H*). Arrowheads point to selected amino acid residues.

### The AlfA monomer structure shows a subdomain deletion and an unusual nucleotide-binding mode

For all previous actin-like filament structures determined by cryo-EM, reliable structures of the monomer obtained by X-ray crystallography were available to guide model building (4, 5, 10). With AlfA, such aid was not at hand, and our efforts to crystallise AlfA yielded only poorly diffracting crystals. However, the cryo-EM map of AlfA was good enough for assisted *de novo* structure solution. The initial model, generated computationally by homology modelling, was manually fitted, adjusted and refined to best explain the cryo-EM density. Model statistics are very good and refinement of the atomic model against the density map was performed both in real and reciprocal space, resulting in a final R-factor of 0.286 (Table S1).

The structure of the monomer reveals the novelty of the particular adaptation of the canonical actin fold, most notably the deletion of an entire subdomain (Fig. 2C). Actins have a butterfly-like domain structure, where the nucleotide-binding pocket is found in a cleft between domains I and II (Fig. 2A). The domains are divided further into four subdomains: IA, IB, IIA and IIB (also numbered 1, 2, 3 and 4), based on the F-actin model (24). The model of AlfA obtained through our cryo-EM reconstruction clearly shows the absence of subdomain IIB, otherwise present in all other known actin-like proteins (Fig. 2A-B), explaining the smaller size of the protein. A structural alignment of representative actin-like protein sequences shows the boundaries and the position of the deletion relative to other proteins (Fig. S4). At sequence level, AlfA is as different from other bacterial actins ParM and MreB as all three are from their eukaryotic counterpart (15). In the remaining subdomains, AlfA shows strongest resemblance to the other DNA-segregating actin-like protein, ParM (Fig. 2B.), especially in the way in which the β-hairpin from domain IB complements the β-barrel-like structure of IA to form a cradle around the central helix of this domain (perpendicular to the plane of Fig. 2A). When we compared the fold of AlfA to all protein structures deposited in the PDB (25), ParM occupied all of the top positions of the list, with the best match having a Z-score of 9.1 and a root-mean-square deviation (RMSD) of the amino acid backbone of 2.34 Å (Fig. S3). Taking into account the 18% sequence identity between ParM and AlfA, this is a striking example of conservation of structure.

**Fig. 2.**
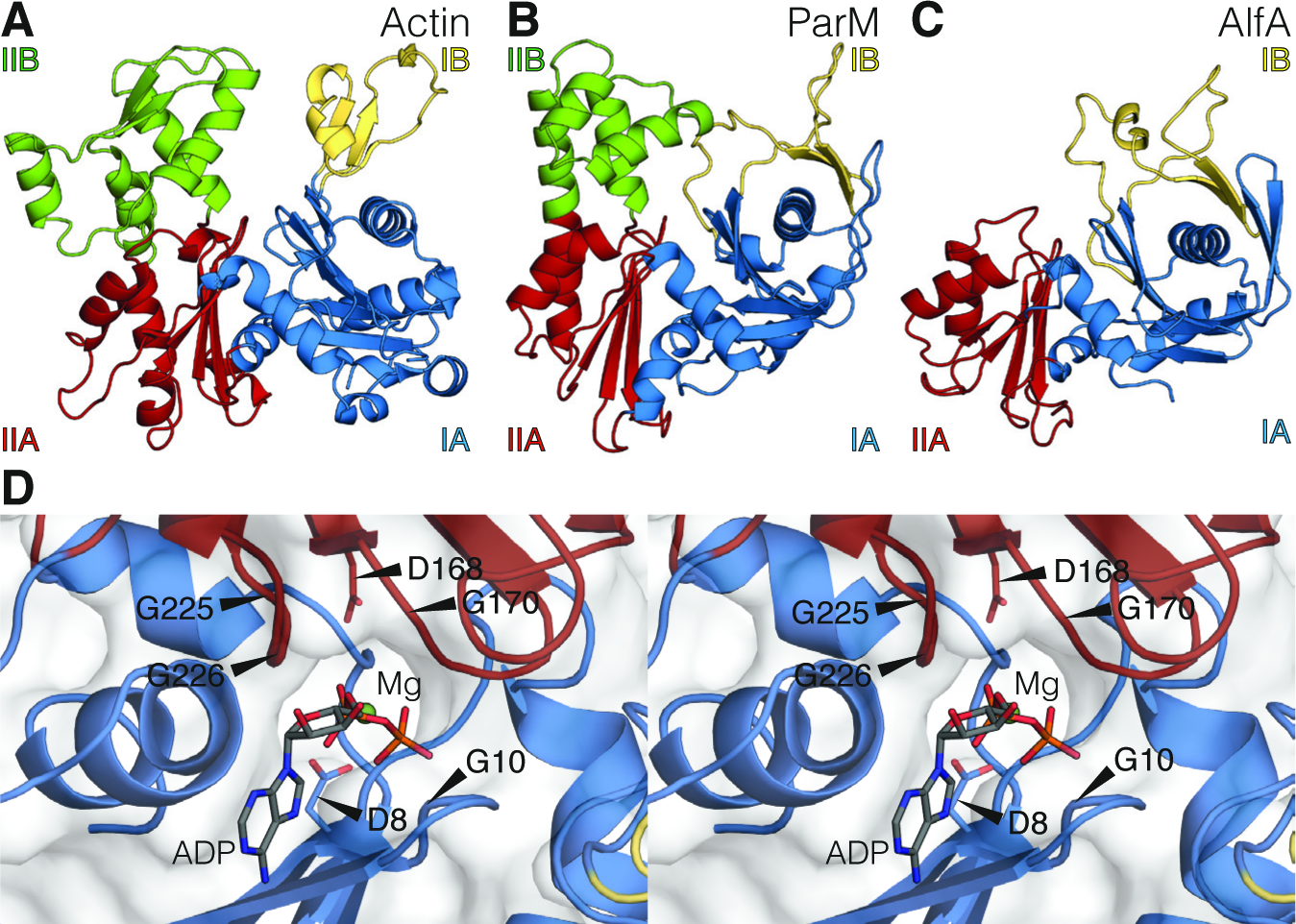
Structural features of AlfA monomer. (*A-C*) Differences in fold and subdomain composition of three actin-like proteins: F-actin from *Oryctolagus cuniculus* (*A*), ParM from *Escherichia coli* plasmid R1 (*B*) and AlfA from *Bacillus subtilis* plasmid pLS32 (*C*). Structural domains of proteins are coloured and labelled according to their homology to subdomains IA (blue), IB (yellow), IIA (red) and IIB (green) (24). Deletion of the entire subdomain IIB in AlfA is clearly evident (*C*), in agreement with amino acid sequence alignments (Fig. S4). Presented models of F-actin and ParM are from PDB depositions 3J8A and 5AEY (in all figures). (*D*) Stereogram of the nucleotide-binding catalytic pocket of AlfA shows ADP molecule surrounded by the universally conserved amino acid residues (indicated by arrowheads, see Fig. S4). The conformation of the bound nucleotide shows the adenosine moiety rotated away from the glycine doublet by around 120°. In all other studied actin-like proteins, this doublet comes in contact with the adenosine, however in AlfA, the moiety is uniquely placed between residues F12 and Y225 (Fig. S5), which are not conserved in other actins (Fig. S4).

AlfA was initially identified from sequences as a member of the actin-like family proteins, due to the presence of signature nucleotide-binding motifs conserved across eukaryotic actin, FtsA, MreB, and ParM (15). The motifs are found in the loops that protrude into the catalytic pocket and include: a pair called *phosphate 1* and *2*, consisting of the consensus DxG sequence in each and involved in the binding and hydrolysis of the γ-phosphate of ATP, as well as the *adenosine* doublet GG important for the positioning of the nucleotide base (26). Our AlfA sequence alignment shows all three signature sequences (Fig. S4) (27) and our structure places them in the context of the nucleotide-binding pocket of the protein (Fig. 2D). The *phosphate 1* motif is composed of residues D8 and G10 of AlfA and *phosphate 2* is D168 and G170. The overall arrangement of the catalytic aspartate residues and the magnesium co-ordination are consistent with the functional requirement for ATP hydrolysis in other actins (2, 8, 24). The cryo-EM density strongly indicates the presence of ADP in the nucleotide-binding pocket (Fig. 1H), suggesting that the majority of the added ATP had been hydrolysed by AlfA before the grid was vitrified.

The catalytic nucleotide-binding pocket of AlfA, although very similar to other actins, shows one striking structural difference. A doublet of glycines called the *adenosine* motif, although conserved in AlfA in the usual position (G225, G226), does not contact the nucleotide base as expected. Instead, the adenosine moiety of ADP is swung out by about 120° from the vicinity of the GG doublet, where it is located in all other actin-like proteins (Fig 2D). In our model, the adenine purine ring of the nucleotide is sandwiched between a pair of aromatic residues F12 and Y255 (Fig. S5), which are not conserved across the actin-like family (Fig. S4). It has been shown that AlfA accepts the other purine nucleotide GTP as a substrate for polymerisation (18) and this can be explained by the fact that the differentially substituted positions of the ATP or GTP purine rings (C2 and C6) are not in contact with the protein. Other actin-like proteins also have the ability to hydrolyse both nucleotides (28, 29), and similarly, most likely not through some shared readout mechanism, but due to a lack of contact between the protein and the substituted positions on the nucleotide.

### Loss of subdomain IIB is accompanied by changes in longitudinal and latitudinal contacts of the AlfA filament

The subdomain deletion found in AlfA poses an interesting question about the maintenance of protofilament (strand) contacts in the polymerised filament. All studied actin-like filament structures show a pair of protofilaments in the native state (Fig. S2). With the notable exception of MreB, the protofilaments wrap around each other in a parallel double helix. For each protein monomer in a double helix, we can define two types of interactions: latitudinal (inter-protofilament) contacts, between monomers situated in the two protofilaments, and longitudinal (intra-protofilament) contacts, between monomers in the same protofilament. In order to assess the architecture of the filament assembly and the consequences of the IIB domain deletion, we visualised the longitudinal (Fig. 3) and latitudinal (Fig. 4) contacts in our AlfA filament model. We knew from previous studies (10), that the contacts are not well conserved on the level of amino acid sequence, so we turned to investigating the internal filament surfaces instead, taking a 4 Å radius of interaction (the upper length limit of a hydrogen bond).

In both actin (Fig. 3A) and ParM (Fig. 3B) the longitudinal contacts are formed through the interactions of the IIB and IB subdomains of one monomer with the IIA and IA subdomains of the other. The interaction of the IB subdomain occurs mainly via the hydrophobic D-loop, which in filamentous actin and ParM inserts into a pocket on the subdomain IA (Fig. 3A-B) in the preceding subunit in the same protofilament. In the case of AlfA, subdomain IIB is absent and so the interaction between the D-loop and the rest of the domain IB is shifted towards the IIA subdomain, with subdomain IB making no contact with the IA subdomain (Fig. 3C). When compared to the other protofilament structures, the remainder of the AlfA monomer in the protofilament is twisted in the direction of where its missing IIB subdomain would be, and the IA subdomain no longer makes any filament contacts, pushed out and away from the filament axis. The longitudinal contacts in the filament thus involve only two domains: IIA and IB, in alternating succession.

**Fig. 3.**
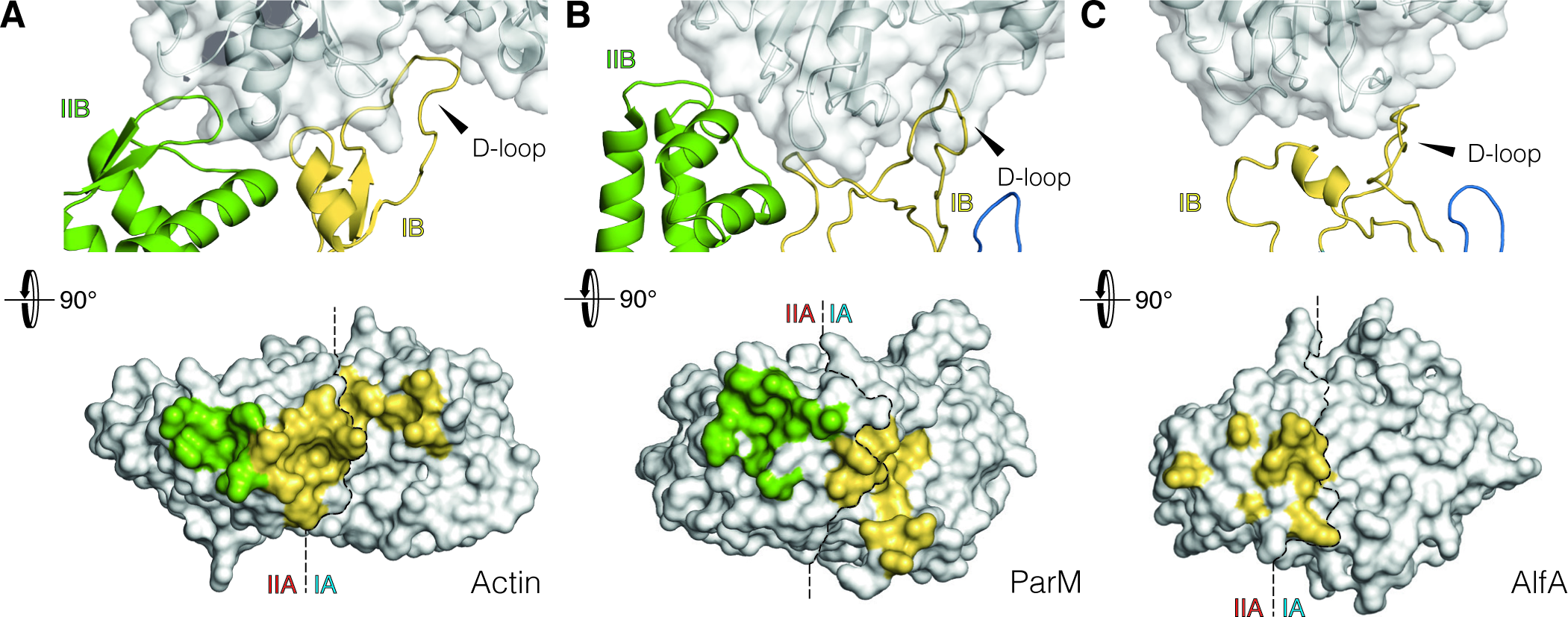
Longitudinal interfaces of actin-like filaments. (*A-C*) Longitudinal (intraprotofilament/strand) interactions in F-actin (*A*), ParM (*B*) and AlfA (*C*). In each of the three examples, the D-loop of the IB subdomain of one monomer in the protofilament makes contact with the monomer placed further up the protofilament. In AlfA (*C*), subdomain IIB, present in ParM (*B*) and F-actin (*A*), as well as other actin-like proteins, is absent and the longitudinal contact is solely via subdomain IB. On the bottom of each panel, the same interaction is shown as an imprint onto the surface of a protofilament monomer, with colours representing amino acid residues in the 4 Å vicinity of the subdomains making the contact. The subdomain divisions of the monomer are indicated by dashed lines. AlfA shows a reduction in surface area and a shift of the IB subdomain interface in comparison to F-actin and ParM.

Actin, ParM and AlfA form staggered filaments, which means that each monomer interacts with two monomers on the opposite protofilament, forming latitudinal contacts that hold the double filaments together. In actin and ParM, all four subdomains make latitudinal contacts with adjacent protofilament monomers (Fig. 4A-B). Notably, the latitudinal contacts of ParM constitute a smaller surface than those of actin (10), although predominantly high-energy salt bridges between long side chains are involved. The structure of AlfA shows a more pronounced departure from the actin filament structure (Fig. 4C). In this case, only the subdomains IB and IIA are responsible for forming the contact, interacting with subdomains IB and IIA of the opposite protofilament.

**Fig. 4.**
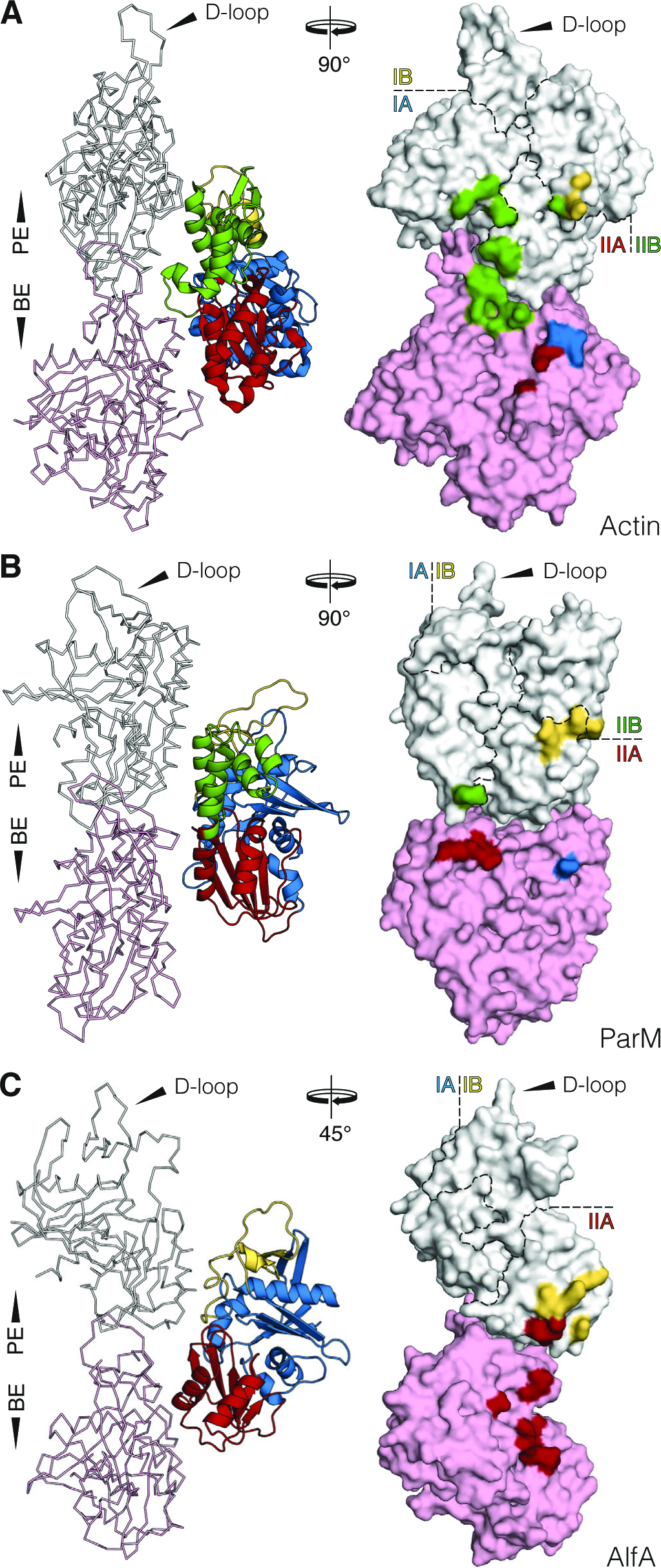
Latitudinal interfaces of actin-like filaments. (*A-C*) Latitudinal (interprotofilament/strand) interactions in F-actin (*A*), ParM (*B*) and AlfA (*C*). On the left side of each panel, three monomers of the filament forming the latitudinal contact are shown, the axis of the filament running down the page. On the right side, interactions are shown as an imprint onto the surface of protofilament monomers, with colours representing amino acid residues in the 4 Å vicinity of the subdomains making the contact. As these filaments are staggered, each interacting monomer is in contact with two monomers in the opposite protofilament. The subdomain divisions of the monomer are indicated by dashed lines. Only subdomains IA and IIB of AlfA are involved in the contact, in alternating order.

**Fig. 5.**
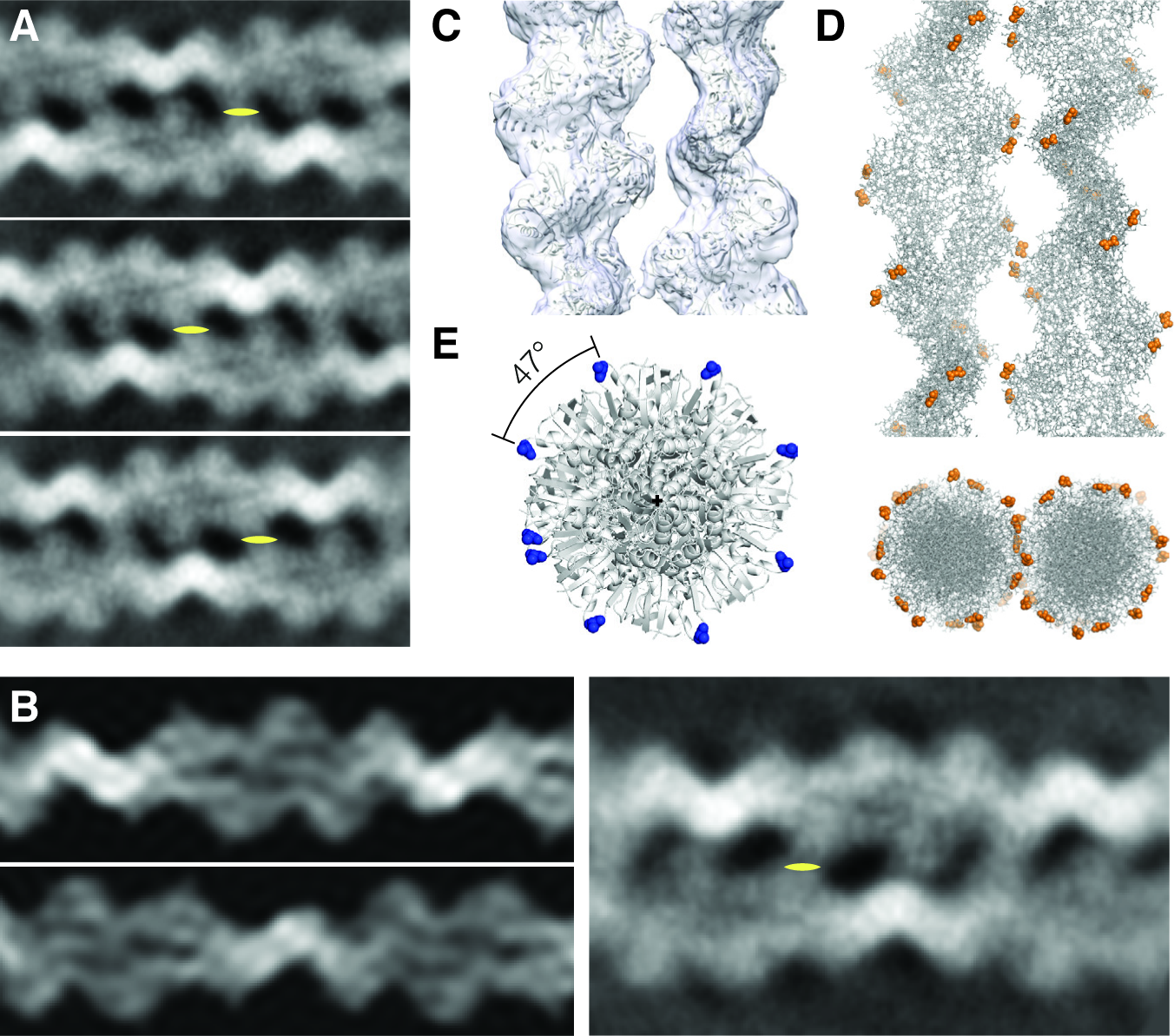
Cryo-EM reconstruction of AlfA filament doublet. (*A*) Representative reference-free 2D class averages of wild-type AlfA show pairs of filaments. The images show two filaments running antiparallel to each other and perfectly out of phase with the narrow part of one filament lining up with the wide part of the other. Yellow symbols indicate the positions of one of the multiple 2-fold symmetry axes perpendicular to the filament axis. (*B*) On the right side of the panel is another reference-free 2D class similar to those in A. On the left side are two simulated projections of the high-resolution filament structure from Fig. 1A, rotated along the filament axis and, in case of the bottom projection, flipped into antiparallel orientation to match the 2D class average on the right. (*C*) Cryo-EM envelope of the AlfA filament doublet, reconstructed from 2D-classified segments using single-particle, non-helical mode of RELION. The filament model has been fitted into the obtained density in the orientation derived from the projection matching in B. (*D*) Side (top) and top view (*bottom*) of the doublet model, in stick representation, with orange spheres indicating the positions of the lysine residues K21, K22, K101 and K102 that stop bundling upon mutation to alanine. (*E*) A closer look down the filament axis with alanine 22 marked in blue, showing the almost perfect 8-fold symmetry of the filament (47° between subunits in one protofilament). We propose that the close-to-integer symmetry enables efficient bundling into doublets.

A quantitative analysis of the protofilament interface differences between the three analysed examples was carried out (Table S2) (30). In case of actin, the longitudinal interactions cover 2299 Å^2^ or 13.9% of the molecule surface. For ParM and AlfA these values are respectively: 1989 Å^2^ (12.6%) and 1051 Å^2^ (8.3%), reflecting the decrease in the interaction surface shown in Fig. 3. Similarly, in the latitudinal contact, 935 Å^2^ or 5.6% of each actin monomer interacts with the two monomers of the opposite protofilament, with the corresponding values for ParM and AlfA being: 746 Å^2^ (4.7%) and 808 Å (6.3%). The apparent increase in latitudinal contact in AlfA is slight; but when we look at the fraction of the total interface of AlfA that is involved in making longitudinal contact, it is 56.5%, compared to 71.1% for actin and 72.7% for ParM. We also see an apparent decrease in the proportion of the total surface of the AlfA monomer involved in interfaces: 14.6%, compared to the 17.3% for ParM and 19.5% for actin. It seems that the loss of subdomain IIB in AlfA places higher emphasis on the latitudinal contacts, but this is perhaps not unexpected. On the whole, AlfA seems to present a further evolutionary departure from eukaryotic actin filament than ParM, not only in terms of amino acid sequence and length, but also the surface implicated in forming filament contacts. This observation is the basis for our notion that AlfA is a simplified actin-like system: despite reductions in size and interaction surfaces, the protein still polymerises into dynamic filaments that bind adaptors and perform mechanical work to fulfil their cellular function.

### Reconstruction of AlfA doublets show antiparallel arrangement necessary for plasmid segregation with a bipolar spindle

Previously published studies (17, 18), indicated that polymerised AlfA forms antiparallel bundles. This property is crucial for the creation of bipolar segregating spindles, as seen in the case of the other DNA-segregating protein ParM (10). The antiparallel orientation of the filaments is necessary to ensure that when new monomers are added to the spindle, it grows in both directions and pushes the plasmid copies apart (9). To study the architecture of the AlfA bundles at a greater level of detail, we decided to use cryo-EM to reconstruct the structure of filament arrangements formed by AlfA.

Taking our cryo-EM micrographs of polymerised, wild-type AlfA (Fig. S1A-D), we obtained reference-free averaged images of pairs of AlfA filaments present in the vitrified sample (Fig. 5A). The images clearly show the two filaments perfectly out of phase, with narrow and thin parts interlocking, as well as some secondary structural features, indicating the quality of the averaging. A close inspection of the images shows the presence of multiple 2-fold axes perpendicular to the plane of the doublet, suggesting that the filaments are antiparallel (rotated by 180°) (Fig. 5A). Indeed, all the obtained averages show filaments in such an arrangement. To determine the angular orientations of the two filaments, we projected and rotated our high resolution AlfA model to match the averages (Fig. 1B). The analysis revealed that the two filaments, although running in opposite directions, have the same orientation, or in other words, are rotated around their main axes to the same angle.

This observation suggests a screw-like and exactly anti-phasic arrangement of the filaments in the spindle, very similar to ParM (10). Using the particles from the 2D classification above, we decided to reconstruct a three-dimensional model of the doublet, with the single-particle (non-helical) mode of RELION. Resolution of the resulting reconstruction was limited to above 10 Å, most likely due to the scarcity of the side views of the filament doublet in the micrographs, but it allowed us to fit the AlfA filament models into the density envelope (Fig 1C). The reconstruction shows the two filaments arranged in the predicted antiparallel fashion, according to angles derived in projection matching (Fig 1D).

Individual monomers in each protofilament of the AlfA filament are related to each other by a left-handed twist of 47° (Fig. 1E). Consequently, when viewed down the main filament axis, the subunits show an almost perfect 8-fold (octagonal) symmetry (Fig *1E*), or integer symmetry. Thus, when two filaments associate into an antiparallel doublet, the inter-filament interactions holding the filaments together are (almost) preserved in each turn of the helix by the symmetries of the filaments. The twist of the high-resolution structure model deviates from the perfect 8-fold symmetry by 2°, but in principle could be adjusted for by a superhelical twist of the doublet and/or conformational variation; indeed, some degree of flexibility is seen in the AlfA filament (Fig. S1). A striking example of this type of doublet symmetrisation is observed in the ParM filament, where the protofilament monomer rotation of 29.8° creates an almost perfect 12-fold symmetry (9, 10).

### Implications of Plasmid Evolution on Structure of AlfA Filament

Both on the level of amino acid sequence, as well as structure, AlfA represents the furthest evolutionary detour from the canonical actin-like protein model. Despite the striking differences in structure described here, AlfA still forms actin-like filaments, which in their function and construction are very close to ParM. Both proteins hydrolyse ATP and polymerise into bipolar spindles and both are capable of segregating plasmid DNA. Moreover, ParM and AlfA have special accessory proteins ParR and AlfB, which bind centromeric sequences of the plasmids they help to separate. Now we are able to show that ParM and AlfA share a common architectural paradigm: maintenance of filament symmetry that ensures the formation of antiparallel, interlocking filament pairs. For AlfA, it is realised through most economical means, which involve a structural subdomain deletion accompanied by an overall reduction of total filament interface area, both requiring a rearrangement of longitudinal and latitudinal surface contacts. The reasons that caused the reduction as well as its evolutionary past are not clear. It may be that it captures particular aspects of plasmid evolution, such as drive for functional simplification and sequence reduction. In such case, AlfA is a perfect example of streamlining functionality and complexity, a distinctive manifestation of a DNA-segregation system, which is able to effectively separate 70-kbp pieces of DNA with just two proteins and some ATP.

## Materials and Methods

### Cloning, expression and purification of wild-type and non-bundling *Bacillus subtilis* AlfA

The AlfA gene from *Bacillus subtilis* subsp. *natto* plasmid pLS32 [National Center for Biotechnology Information (NCBI) Database ID code WP_013603336.1] was codon-optimised for expression in *E. coli* and synthesised (IDT Inc.). The gene was cloned by PCR and Gibson assembly into expression vector pOPTL using Q5 DNA polymerase and Gibson Assembly Master Mix (New England Biolabs Inc.). Protein production was carried out in *E. coli* strain C41(DE3) (Lucigen Corp.). Cloning into the vector created an N-terminal fusion of AlfA with a poly-histidine tagged lipoyl domain from *Bacillus stearothermophilus* pyruvate dehydrogenase complex, helping achieve high protein yields; the vector also possesses a TEV protease cleavage site between the target and the fusion, which leaves GSH residues ahead of the N-terminal methionine after TEV protease cleavage. Six litres of 2× TY media supplemented with 100 μg/mL ampicillin were directly inoculated with the bacterial lawns collected from six agar plates grown overnight. The culture was incubated at 37 °C with shaking at 200 rpm. When OD600 of 1.0 was reached, protein expression was induced with 500 μM isopropyl β-D-1-thiogalactopyranoside (IPTG). The culture was incubated for a further 15 h at 20 °C and harvested. The entire cell pellet was sonicated on ice after resuspension in 200 mL of NiA buffer [50 mM Tris˙HCl, 300 mM KCl, 2 mM tris(2-carboxyethyl)phosphine (TCEP), 1 mM NaN3, pH 8.0] supplemented with some DNAse I, RNase E and lysozyme. The lysate was cleared of insoluble debris by centrifugation in a Ti 45 rotor (Beckman-Coulter Inc.) at 45,000 rpm and 4 °C for 45 min. Clarified lysate was mixed with 20 mL of washed Ni-NTA Agarose beads (Qiagen N.V.) and incubated for 1 h at 4 °C with gentle agitation. The beads were washed with 400 mL NiA buffer mixed with 2% (vol/vol) NiB buffer (50 mM Tris˙HCl, 300 mM KCl, 1 M imidazole, 2 mM TCEP, 1mM NaN3, pH 8.0). The beads were suspended in 30 mL of NiA buffer, 500 μg of TEV protease (Sigma-Aldrich Corp.) were added to the suspension and incubated overnight at 4 °C. The supernatant was collected, the beads were washed with further 30 mL of NiA buffer and the two obtained fractions were pooled and concentrated in Amicon Ultra-15 centrifugal filter unit [10 kDa molecular weight cut-off (MWCO); Merck KGaA]. The concentrate was separated on HiPrep Sephacryl S-300 16/60 size-exclusion chromatography column (GE Healthcare Corp.) equilibrated with TENT100 buffer (20 mM Tris·HCl, 100 mM KCl, 2 mM TCEP, 1 mM EDTA, 1mM NaN3, pH 8.0). The protein eluted as a single peak, and the corresponding fractions were pooled and concentrated as above. Presence of a low-molecular-weight peak with high absorbance at 260 nm suggested the removal of the co-purified and natively bound nucleotide. Purity of the protein preparation was confirmed by SDS-PAGE, and the protein was flash-frozen in liquid nitrogen in 100 μL aliquots at 20 mg/mL. The non-bundling mutant of AlfA was cloned, expressed and purified in the same manner as the wild-type protein.

### Cryo-EM sample preparation and data collection for AlfA filament and doublet structure reconstruction

For structure solution with cryo-EM, concentrated AlfA protein solution was diluted from 20 mg/mL to 0.5 mg/mL in 20 mM Tris·HCl, 100 mM KCl, 2 mM TCEP, 1mM NaN3, pH 8.0. To polymerise AlfA, 5 mM ATP and 10 mM MgCl_2_ were added to the diluted protein solution and used immediately. 3 μL of the solution were applied onto a Cu 300 mesh EM grid with R 1.2/1.3 Quantifoil holey carbon support film (Quantifoil Micro Tools GmbH), that had been glow-discharged prior to use. The sample on the grid was vitrified in liquid ethane at below −160 °C using a Vitrobot Mark IV (FEI Comp.). The cryo-EM grids prepared with non-bundling AlfA mutant were imaged using Titan Krios G3 transmission electron microscope (FEI Comp.) at 300 kV accelerating voltage and liquid nitrogen temperature. The images were recorded using automated data acquisition software EPU (FEI Comp.) on a Falcon 3EC direct electron detector (FEI Comp.) operated in electron-counting mode. Each exposure lasted 60 s and was collected as a 75-frame movie using a total electron dose of 33 e^−^/A^2^ at 1.07 Å pixel size and defocus values between –2.7 and –1.5 μm.

For doublet structure solution with wild-type AlfA, the samples were prepared as above using Cu/Rh 300 mesh EM grid with R 2/2 Quantifoil holey carbon support film (Quantifoil Micro Tools GmbH) and imaged with Tecnai Polara G2 transmission electron microscope (FEI Comp.) at 300 kV accelerating voltage and liquid nitrogen temperature. The images were recorded on Falcon III direct electron detector (FEI Comp.) operated in integrating mode. Each exposure lasted 1.5 s and was collected as a 46-frame movie using a total electron dose of 39 e^−^/A^2^ at 1.34 Å pixel size and defocus values between –2.7 and –1.5 μm.

### Cryo-EM data processing for AlfA filament structure solution

In total, 818 movies were acquired during a single 24-hour data collection session. The dataset was processed in RELION 2.0 (20) using the helical reconstruction method (21). The images were corrected for beam-induced motion with MotionCor2 (31) using 5 × 5 patches and dose-weighting and contrast transfer functions (CTFs) were estimated with Gctf (32) using nondose-weighted images. From the dataset, 386 movies with estimated resolutions below 4.0 Å were selected for further processing. A few hundred filament segments were picked manually and extracted in 350-pixel boxes for 2D classification. Good classes were used as templates for automated particle picking, which was optimised for efficient selection of AlfA single filaments. Auto-picked particles were then extracted as helical segments and 2D-classified as such. In total, 78,786 best particles (helical segments) from 30 2D classes were selected and used for 3D auto-refinement. For this step a reference was created from a ParM monomer crystal structure (PDB: 1MWM), which was extended into a helix, simulated as a density map, and filtered to 30 Å. Initial helical parameters used for reference creation and 3D auto-refinement were derived from an analysis of the 2D classes, as well as previously published data (18). The initial 3D auto-refinement was followed by movie refinement and particle polishing (33), yielding particles with a higher signal-to-noise ratio, used subsequently in a final round of 3D auto-refinement. For this round, a solvent mask covering 30% of the central helix Z-length was used to calculate solvent-flattened FSC curves in each refinement step. The obtained map was post-processed with a mask covering 20% of the Z-length and a raised-cosine soft edge of 8 pixels. The final overall and local resolution were assessed by the gold-standard FSC procedure implemented in RELION, using the FSC0.143 criterion (22).

For the purpose of model building, an initial AlfA monomer model was created using the SWISS-MODEL server (34). The model was fitted into a central portion of the 3D refined and post-processed cryo-EM map, cut out using REFMAC (35) and corresponding to a single filament subunit. The model was manually adjusted with MAIN (36) and refined in reciprocal space against the cryo-EM density using REFMAC (37) and in real space using PHENIX (38). To allow refinement in reciprocal space, the cryo-EM map was back-transformed into structure factors (REFMAC in SFCALC mode). The fitted and refined map was assessed using the standard crystallographic R-factor analysis and the model was judged for stereochemical plausibility with MolProbity (39).

### Cryo-EM data processing for AlfA doublet reconstruction

In total, 976 movies were acquired during a single 24-hour data collection session. The dataset was corrected for beam-induced motion and CTF as above, with 717 images selected for further processing in RELION 2.0. A few hundred filament doublets were manually picked and extracted as single particles (not helical segments) in 300-pixel boxes. These were 2D-classified and used as references for auto-picking. The set of auto-picked particles was 2D-classified again and 68,986 final particles were selected for 3D auto-refinement using a filament doublet model prepared from the AlfA filament structure and filtered to 60 Å resolution. The obtained map was post-processed using a solvent mask with raised-cosine soft edge of 8 pixels.

For the projection matching, the *relion_project* utility of RELION was used to generate a set of projections of the AlfA filament rotated at 5° increments around the main axis. From the set, two projections matching a representative 2D class average were manually selected. The rotational angles derived from the selection were applied to AlfA single filament model and fitted into the obtained density to reconstruct the filament doublet.

### Structure analysis of AlfA and other actin-like filaments

Amino acid sequence alignment of actin-like proteins guided by secondary structure elements was carried out using PROMALS3D (27). The analysis of filament interfaces of actin, ParM and AlfA was conducted with PDBe PISA 1.52 (30). The structural similarity search was carried out using PDBe FOLD 2.59 (25).

### Coordinates and map Depositions

The refined filament structure was deposited in the PDB with ID code XXXX, and the corresponding cryo-EM 3D map was deposited in the EMDataBank (EMDB) with ID code: EMD-XXXX.

---

**Fig. S1.**
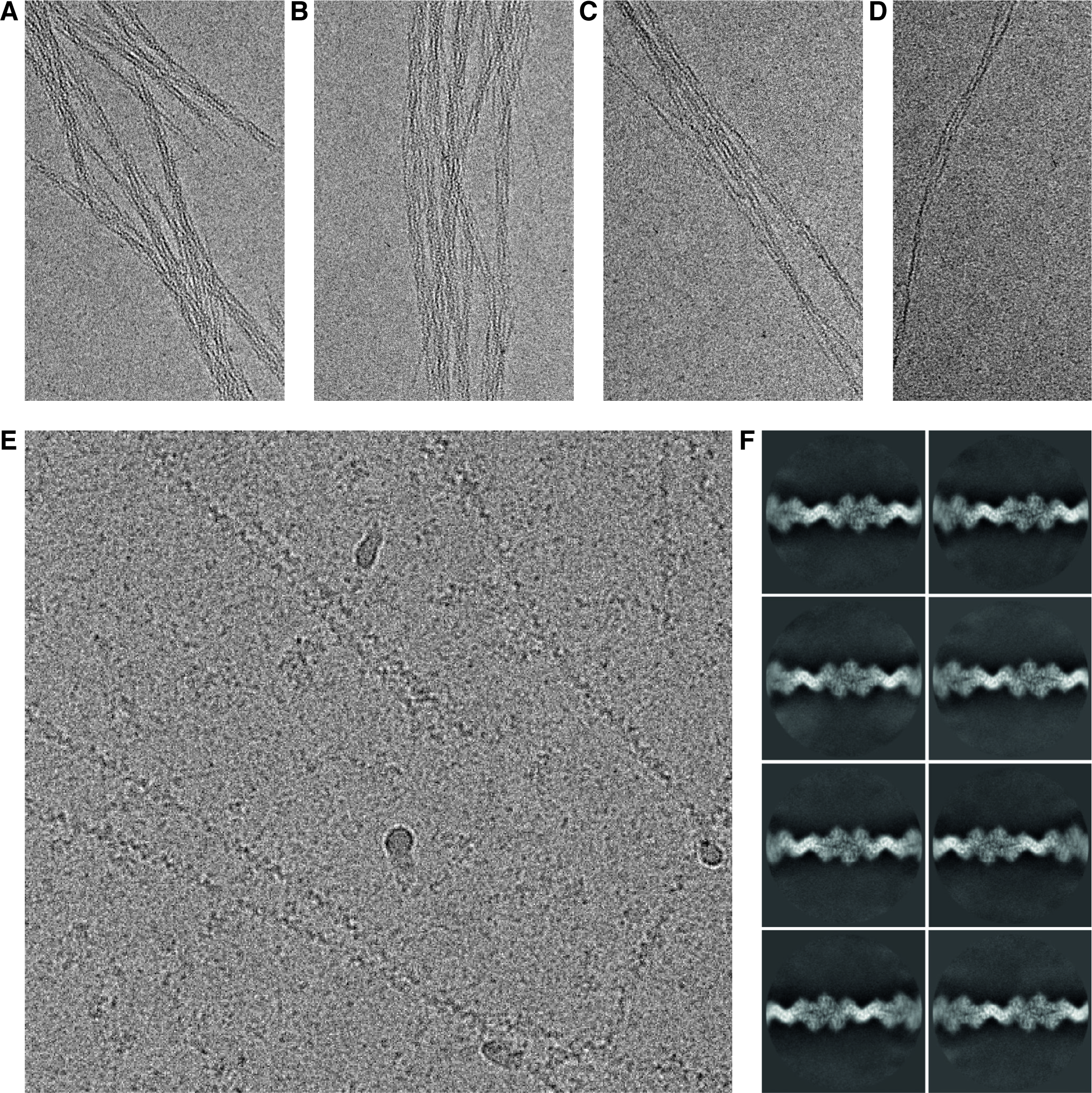
Electron cryo-microscopic imaging of AlfA filaments. (A-D) Images show polymerised wild-type AlfA in vitrified ice. The filaments gather into bundles that disband with increasing salt (KCl) concentration: 50 mM (*A*), 200 mM (*B*), 500 mM (*C*) and 1000 mM (*D*). The micrographs were taken on a FEI Tecnai T12 microscope at 120 kV. (*E*) Polymerised, non-bundling AlfA mutant shown forming single filaments with 50 mM KCl. This image was taken on a FEI Titan Krios G3 microscope at 300 kV, on a Falcon 3 camera in electron counting mode, as part of the dataset used for the high-resolution structure solution (Fig. 1). (*F*) Reference-free averages of filament segments from the dataset, obtained in RELION (*X*) via 2D classification. The images show AlfA filaments composed of protofilaments arranged in a double helix with staggered subunits. For all microscopy purposes, 0.5 mg/mL of purified AlfA was used. The remaining buffer ingredients were: 20 mM Tris·HCl (pH 8.0), 2 mM TCEP and 1mM NaN3. Protein filaments were polymerised via the addition of 5 mM ATP and 10 mM MgCl_2_.

**Fig. S2.**
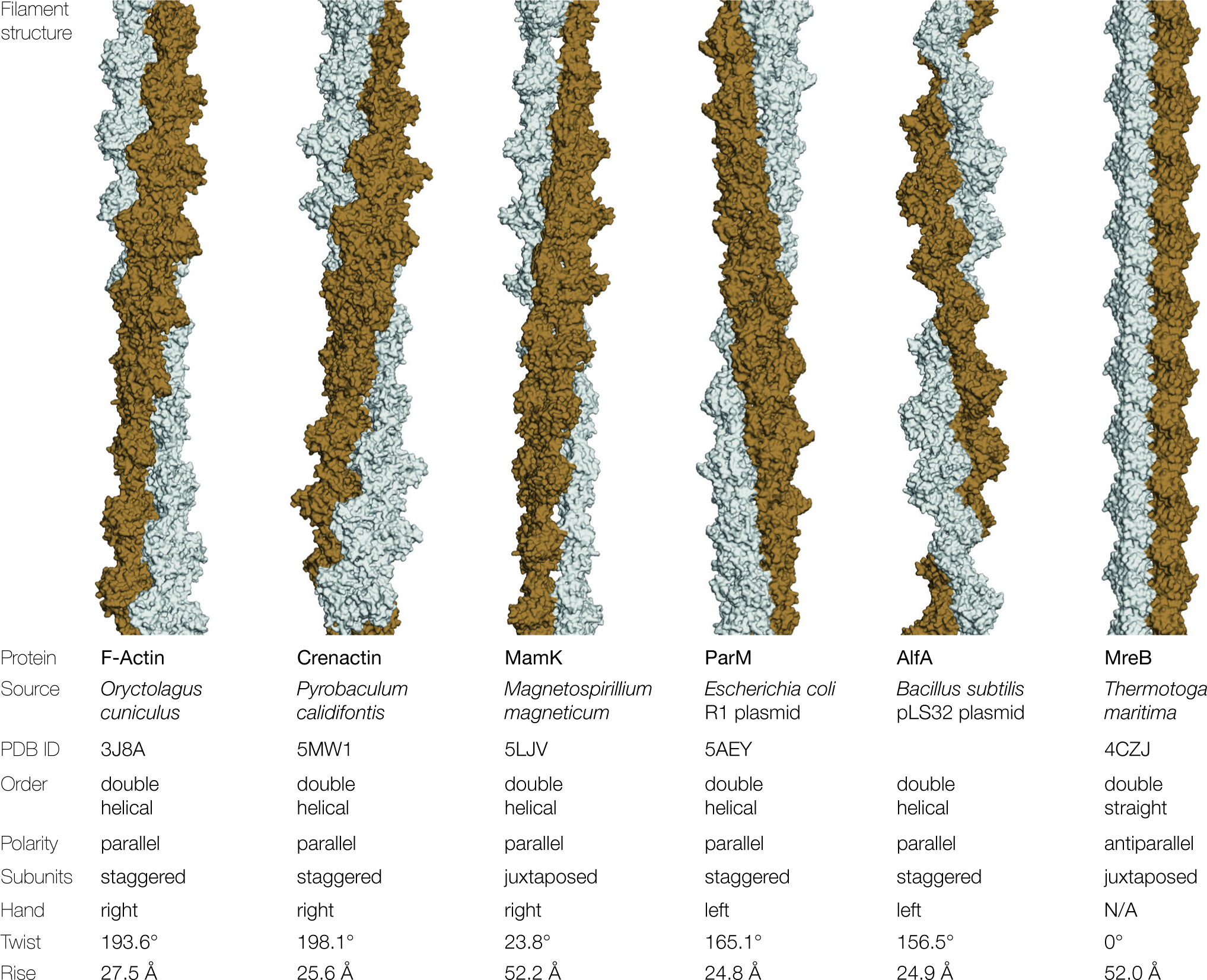
Architecture of actin-like double filaments. Comparison of known atomic structures of double filaments formed by F-actin and actin-like proteins crenactin, MamK, ParM, AlfA and MreB. Each filament is composed of a parallel doublet of protofilaments twisted into a double helix, apart from MreB, which forms an antiparallel, straight filament. Source organism, PDB ID, order, polarity and subunit arrangement are indicated, together with the filament's helical parameters. The architecture of AlfA is closest to ParM, but the protofilaments are more twisted and narrower at the crossover. Note that the height of the subunit in the filament, which is twice the value of the helical rise for staggered filaments and equal to the rise for juxtaposed filaments, is very similar across all studied actin-like filaments, including the smaller AlfA.

**Fig. S3.**
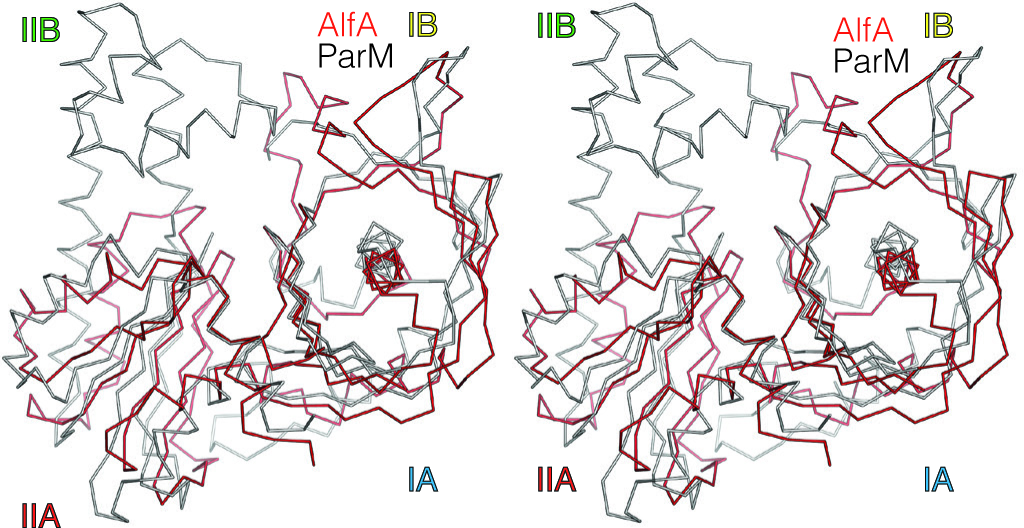
Structural similarity between ParM and AlfA monomers. Stereogram showing a superposition of amino acid backbone (Cα) trace of AlfA (red) and ParM (grey) monomers. The two structures are strikingly similar in fold, despite only 18% sequence identity. The structure of ParM (PDB ID: 4A61) and the alignment shown here are the top result from a PDB-wide search for structural similarity carried out with PDBe FOLD (25). The two agree with each other with a Z-score of 9.1 and RMSD between the C_α_ backbone atoms of 2.34 Å over 230 residues, covering 83% of AlfA and 72% of ParM amino acid sequence.

**Fig. S4.**
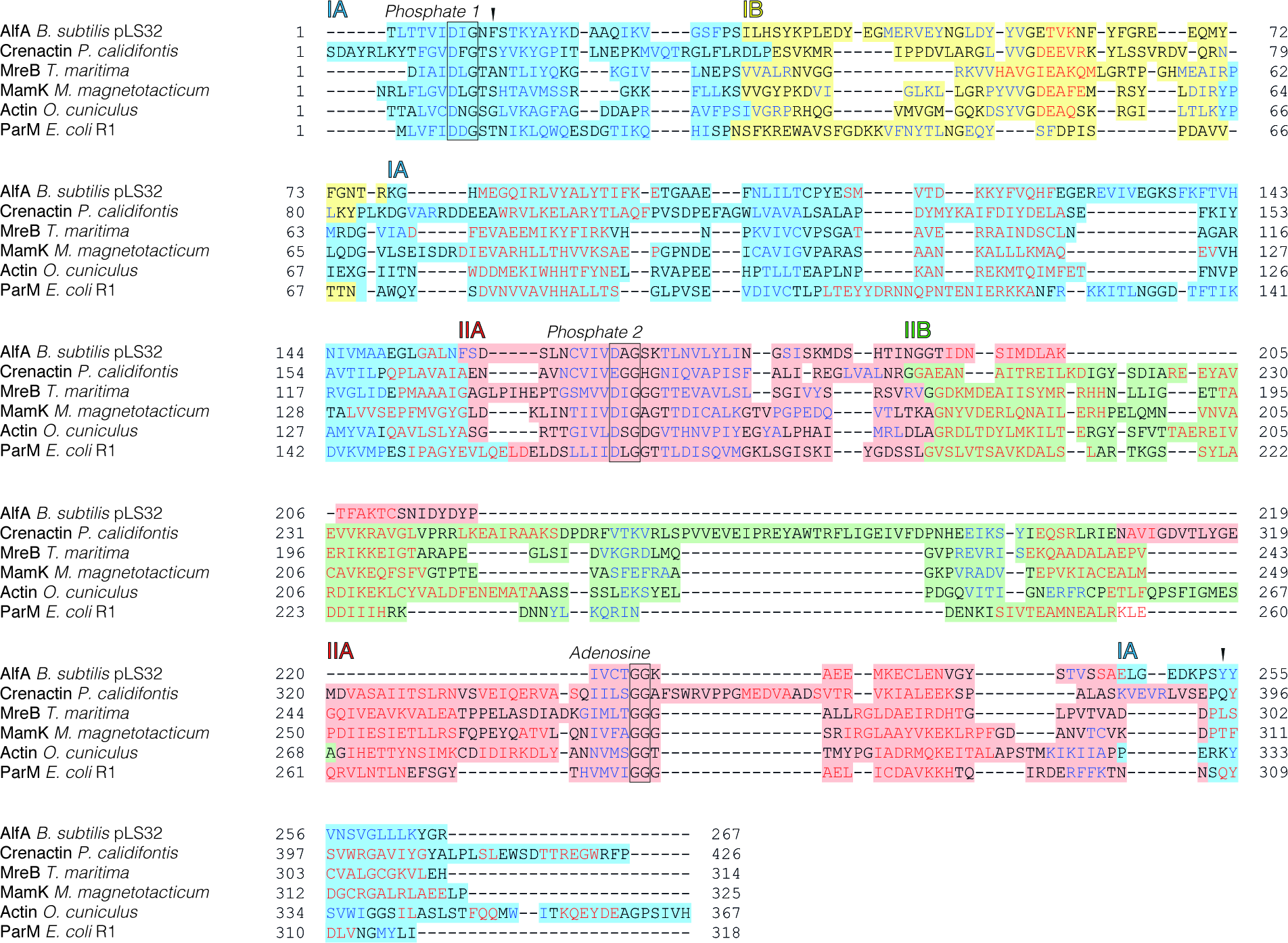
Sequence and secondary structure conservation in actin-like proteins. Shown here is an alignment of amino acid sequences of AlfA, F-actin and other actin-like proteins: crenactin, MamK, ParM and MreB. The alignment was prepared using PROMALS3D (27) and shows divergence and conservation of sequences and secondary structure elements, based on the protein structure models. Red letters indicate α-helices and β-sheet stretches are marked blue. Colour shadings demarcate boundaries between the subdomains: IA (blue), IB (yellow), IIA (red) and IIB (green). Boxed areas indicate positions of universally conserved sequence motifs involved in nucleotide hydrolysis and binding (24). With the deletion of subdomain IIB and novel nucleotide binding mode (Fig. S5), AlfA represents a clear departure from other actin-like proteins. Arrowheads indicate the location of phenylalanine 12 and tryptophan 255, which position the adenine base of ADP in filamentous AlfA. Neither the residues, nor the interaction, are conserved in other studied actin-like proteins.

**Fig. S5.**
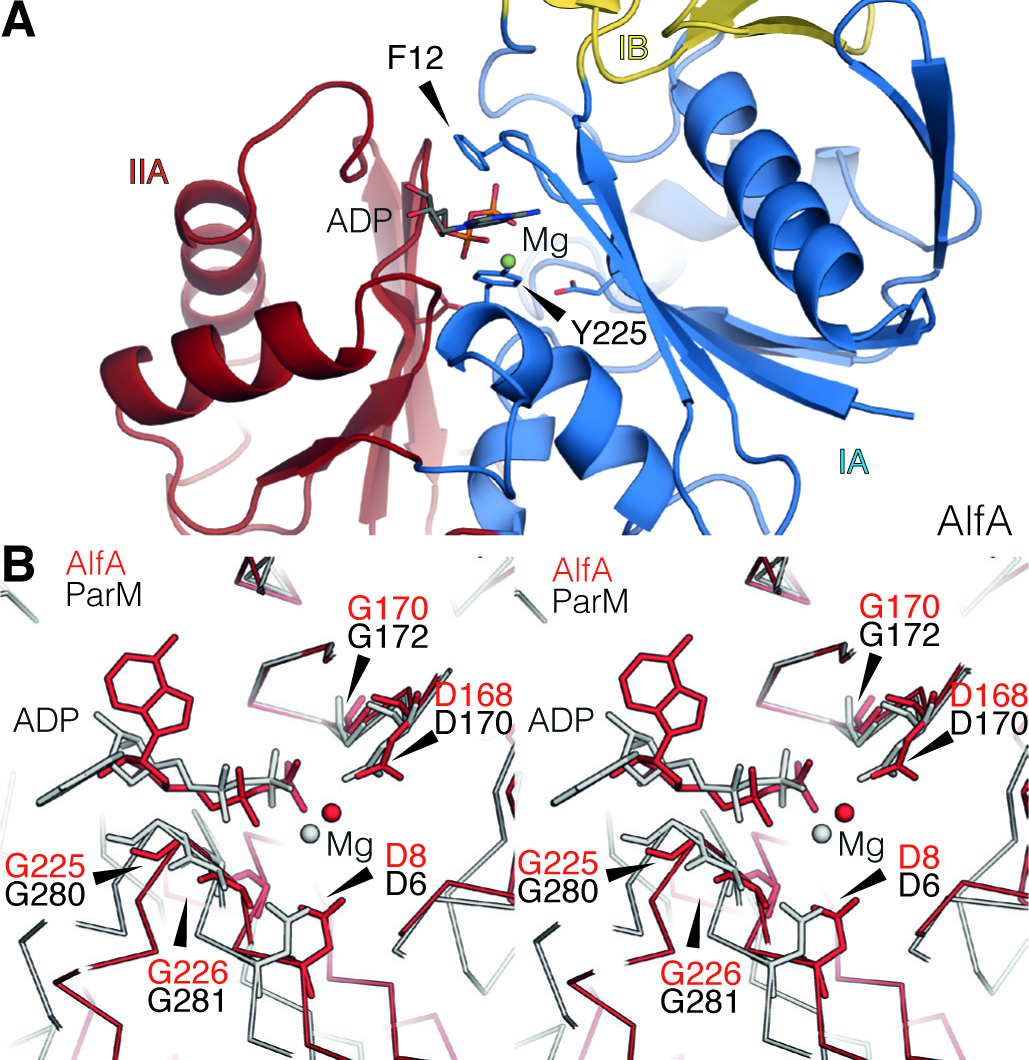
Novel nucleotide binding mode in AlfA. (*A*) In the reconstructed atomic structure of the AlfA filament, the adenosine moiety of ADP molecule is sandwiched between two aromatic residues, Y12 and W255, shown here as skeletal models in a ribbon diagram of the filament monomer. (*B*) Stereogram of amino acid backbone (Cα) trace of AlfA (red) and ParM (grey) monomers in the polymerised filament, showing the nucleotide binding mode and the universally conserved residues involved in the interaction. The adenosine ring and the ribose sugar are rotated away from the glycines 225 and 226, which come in contact with these moieties in ParM and other actin-like proteins.

**Supplementary Table S1.**
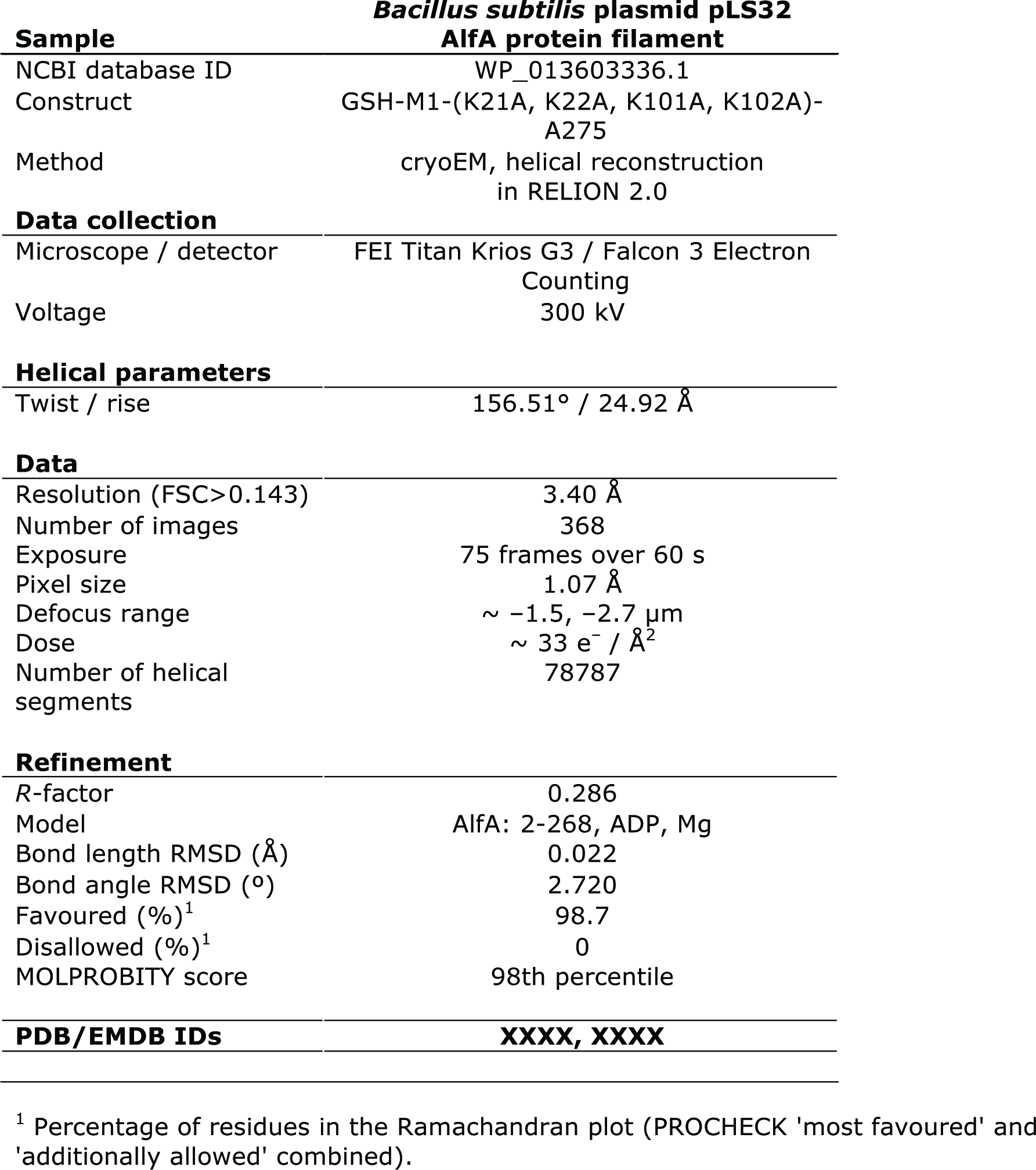
CryoEM and model refinement data.

**Supplementary Table S2.** Surface area analysis of filament interfaces in F-actin, ParM and AlfA.

| **F-Actin** (*O. cuniculus*) | Area (Å<sup>2</sup>) | per tot. surf.<br>(%) | per tot. int.<br>(%) |
| --- | --- | --- | --- |
| 3J8A |  |  |  |
| Monomer surface (total) | 16560.0 |  |  |
| Filament interface (total) | 3234.1 | 19.5 |  |
| Longitudinal | 2299.1 | 13.9 | 71.1 |
| Latitudinal | 935.0 | 5.6 | 28.9 |
| **ParM** (*E. coli*) |  |  |  |
| 5AEY |  |  |  |
| Monomer surface (total) | 15714.0 |  |  |
| Filament interface (total) | 2735.7 | 17.4 |  |
| Longitudinal | 1989.2 | 12.6 | 72.7 |
| Latitudinal | 746.5 | 4.7 | 27.3 |
| **AlfA** (*B. subtilis*) |  |  |  |
| XXXX |  |  |  |
| Monomer surface (total) | 12679.0 |  |  |
| Filament interface (total) | 1859.0 | 14.7 |  |
| Longitudinal | 1051.1 | 8.3 | 56.5 |
| Latitudinal | 807.9 | 6.4 | 43.5 |

## References

1. Izoré T, van den Ent F (2017) Bacterial Actins. Prokaryotic Cytoskeletons, Subcellular Biochemistry. (Springer, Cham), pp 245-266.

2. van den Ent F, Amos LA, Löwe J (2001) Prokaryotic origin of the actin cytoskeleton. Nature 413(6851):39-44.

3. Szwedziak P, Wang Q, Freund SMV, Löwe J (2012) FtsA forms actin-like protofilaments. EMBO J 31(10):2249-2260.

4. Löwe J, He S, Scheres SHW, Savva CG (2016) X-ray and cryo-EM structures of monomeric and filamentous actin-like protein MamK reveal changes associated with polymerization. Proc Natl Acad Sci 113(47):13396-13401.

5. Izoré T, Kureisaite-Ciziene D, McLaughlin SH, Löwe J Crenactin forms actin-like double helical filaments regulated by arcadin-2. eLife 5. doi:10.7554/eLife.21600.

6. Gerdes K, Howard M, Szardenings F (2010) Pushing and Pulling in Prokaryotic DNA Segregation. Cell 141(6):927-942.

7. Jensen RB, Gerdes K (1997) Partitioning of plasmid R1. The ParM protein exhibits ATPase activity and interacts with the centromere-like ParR-parC complex11 Edited by M. Gottesman. J Mol Biol 269(4):505-513.

8. van den Ent F, Moller-Jensen J, Amos LA, Gerdes K, Löwe J (2002) F-actin-like filaments formed by plasmid segregation protein ParM. EMBO J 21(24):6935-6943.

9. Gayathri P, et al. (2012) A Bipolar Spindle of Antiparallel ParM Filaments Drives Bacterial Plasmid Segregation. Science 338(6112):1334-1337.

10. Bharat TAM, Murshudov GN, Sachse C, Löwe J (2015) Structures of actin-like ParM filaments show architecture of plasmid-segregating spindles. Nature 523(7558):106-110.

11. Møller-Jensen J, Ringgaard S, Mercogliano CP, Gerdes K, Löwe J (2007) Structural analysis of the ParR/parC plasmid partition complex. EMBO J 26(20):4413-4422.

12. Tanaka T, Ogura M (1998) A novel Bacillus natto plasmid pLS32 capable of replication in Bacillus subtilis. FEBS Lett 422(2):243-246.

13. Tanaka T (2010) Functional Analysis of the Stability Determinant AlfB of pBET131, a Miniplasmid Derivative of Bacillus subtilis (natto) Plasmid pLS32. J Bacteriol 192(5):1221-1230.

14. Tanaka T, Koshikawa T (1977) Isolation and characterization of four types of plasmids from Bacillus subtilis (natto). J Bacteriol 131(2):699-701.

15. Becker E, et al. (2006) DNA segregation by the bacterial actin AlfA during Bacillus subtilis growth and development. EMBO J 25(24):5919-5931.

16. Tanaka T, Ishida H, Maehara T (2005) Characterization of the Replication Region of Plasmid pLS32 from the Natto Strain of Bacillus subtilis. J Bacteriol 187(13):4315-4326.

17. Polka JK, Kollman JM, Mullins RD (2014) Accessory factors promote AlfA-dependent plasmid segregation by regulating filament nucleation, disassembly, and bundling. Proc Natl Acad Sci 111(6):2176-2181.

18. Polka JK, Kollman JM, Agard DA, Mullins RD (2009) The Structure and Assembly Dynamics of Plasmid Actin AlfA Imply a Novel Mechanism of DNA Segregation. J Bacteriol 191(20):6219-6230.

19. von der Ecken J, et al. (2015) Structure of the F-actin-tropomyosin complex. Nature 519(7541):114-117.

20. Scheres SHW (2012) RELION: Implementation of a Bayesian approach to cryo-EM structure determination. J Struct Biol 180(3):519-530.

21. He S, Scheres SHW (2017) Helical reconstruction in RELION. J Struct Biol 198(3):163-176.

22. Rosenthal PB, Henderson R (2003) Optimal Determination of Particle Orientation, Absolute Hand, and Contrast Loss in Single-particle Electron Cryomicroscopy. J Mol Biol 333(4):721-745.

23. Ghosal D, Löwe J (2015) Collaborative protein filaments. EMBOJ 34(18):2312-2320.

24. Kabsch W, Holmes KC (1995) The actin fold. FASEB J 9(2):167-174.

25. Krissinel E, Henrick K (2004) Secondary-structure matching (SSM), a new tool for fast protein structure alignment in three dimensions. Acta Crystallogr D Biol Crystallogr 60(12): 2256-2268.

26. Bork P, Sander C, Valencia A (1992) An ATPase domain common to prokaryotic cell cycle proteins, sugar kinases, actin, and hsp70 heat shock proteins. Proc Natl Acad Sci U S A 89(16):7290-7294.

27. Pei J, Kim B-H, Grishin NV (2008) PROMALS3D: a tool for multiple protein sequence and structure alignments. Nucleic Acids Res 36(7):2295-2300.

28. Popp D, et al. (2010) Filament structure, organization, and dynamics in MreB sheets. J Biol Chem 285(21):15858-15865.

29. Popp D, et al. (2008) Molecular structure of the ParM polymer and the mechanism leading to its nucleotide-driven dynamic instability. EMBO J 27(3):570-579.

30. Krissinel E, Henrick K (2007) Inference of Macromolecular Assemblies from Crystalline State. J Mol Biol 372(3):774-797.

31. Zheng SQ, et al. (2017) MotionCor2: anisotropic correction of beam-induced motion for improved cryo-electron microscopy. Nat Methods 14(4):331-332.

32. Zhang K (2016) Gctf: Real-time CTF determination and correction. J Struct Biol 193(1):1-12.

33. Scheres SH (2014) Beam-induced motion correction for sub-megadalton cryo-EM particles. eLife 3:e03665.

34. Schwede T, Kopp J, Guex N, Peitsch MC (2003) SWISS-MODEL: an automated protein homology-modeling server. Nucleic Acids Res 31(13):3381-3385.

35. Brown A, et al. (2015) Tools for macromolecular model building and refinement into electron cryo-microscopy reconstructions. Acta Crystallogr D Biol Crystallogr 71(1):136-153.

36. Turk D (2013) MAIN software for density averaging, model building, structure refinement and validation. Acta Crystallogr D Biol Crystallogr 69(Pt 8):1342-1357.

37. Murshudov GN, Vagin AA, Dodson EJ (1997) Refinement of Macromolecular Structures by the Maximum-Likelihood Method. Acta Crystallogr D Biol Crystallogr 53(3):240-255.

38. Adams PD, et al. (2010) PHENIX: a comprehensive Python-based system for macromolecular structure solution. Acta Crystallogr D Biol Crystallogr 66(Pt 2):213-221.

39. Chen VB, et al. (2010) MolProbity: all-atom structure validation for macromolecular crystallography. Acta Crystallogr D Biol Crystallogr 66(1):12-21.

